# Inducible protein degradation as a strategy to identify Phosphoprotein Phosphatase 6 substrates in RAS-mutant colorectal cancer cells

**DOI:** 10.1101/2023.03.25.534211

**Authors:** Natasha C. Mariano, Scott F. Rusin, Isha Nasa, Arminja N. Kettenbach

## Abstract

Protein phosphorylation is an essential regulatory mechanism that controls most cellular processes, including cell cycle progression, cell division, and response to extracellular stimuli, among many others, and is deregulated in many diseases. Protein phosphorylation is coordinated by the opposing activities of protein kinases and protein phosphatases. In eukaryotic cells, most serine/threonine phosphorylation sites are dephosphorylated by members of the Phosphoprotein Phosphatase (PPP) family. However, we only know for a few phosphorylation sites which specific PPP dephosphorylates them. Although natural compounds such as calyculin A and okadaic acid inhibit PPPs at low nanomolar concentrations, no selective chemical PPP inhibitors exist. Here, we demonstrate the utility of endogenous tagging of genomic loci with an auxin-inducible degron (AID) as a strategy to investigate specific PPP signaling. Using Protein Phosphatase 6 (PP6) as an example, we demonstrate how rapidly inducible protein degradation can be employed to identify dephosphorylation SITES and elucidate PP6 biology. Using genome editing, we introduce AID-tags into each allele of the PP6 catalytic subunit (PP6c) in DLD-1 cells expressing the auxin receptor Tir1. Upon rapid auxin-induced degradation of PP6c, we perform quantitative mass spectrometry-based proteomics and phosphoproteomics to identify PP6 substrates in mitosis. PP6 is an essential enzyme with conserved roles in mitosis and growth signaling. Consistently, we identify candidate PP6c-dependent phosphorylation sites on proteins implicated in coordinating the mitotic cell cycle, cytoskeleton, gene expression, and mitogen-activated protein kinase (MAPK) and Hippo signaling. Finally, we demonstrate that PP6c opposes the activation of large tumor suppressor 1 (LATS1) by dephosphorylating Threonine 35 (T35) on Mps One Binder (MOB1), thereby blocking the interaction of MOB1 and LATS1. Our analyses highlight the utility of combining genome engineering, inducible degradation, and multiplexed phosphoproteomics to investigate signaling by individual PPPs on a global level, which is currently limited by the lack of tools for specific interrogation.

## Introduction

Protein phosphorylation is an essential regulatory process that governs most cellular processes, including cell cycle progression, cell division, and response to extracellular stimuli, among many others, and it is deregulated in many diseases. The opposing activities of protein kinases and protein phosphatases determine the phosphorylation state of proteins. In eukaryotic cells, most proteins are phosphorylated on at least one serine, threonine, or tyrosine residue at some point in their lifetime (1). Although hundreds of thousands of phosphorylation sites have been identified, less than 10% have been assigned to a specific kinase and less than 1% to a specific phosphatase (2, 3). Phosphoprotein Phosphatases (PPPs) are highly conserved enzymes that control over 90% of serine/threonine dephosphorylation. In humans, the PPP family includes PP1, PP2A, PP2B, PP4, PP5, PP6, and PP7, which consist of a catalytic subunit that binds additional regulatory subunits (4, 5). Each catalytic subunit contains a highly conserved catalytic domain, and the active site residues are a 100% conserved group of three aspartic acids, two histidines, and an asparagine that coordinate bivalent cations (6). The conserved and shallow nature of the active site and its preference for negative charges makes selectively targeting difficult (6, 7). Naturally occurring toxins, such as calyculin A, microcystin, and okadaic acid, inhibit most PPPs at nanomolar concentrations; however, they lack selectivity (8). Despite their conserved active site, PPPs are highly specific enzymes. They achieve substrate specificity through the formation of dimeric or trimeric holoenzymes and recognition of substrates through short-linear motifs (SLiMs). For instance, PP1 forms dimers and the catalytic subunit recognizes SLiMs in the regulatory subunits, which are often substrates (9, 10). In trimeric holoenzymes, such as PP2A and PP4, the regulatory subunits recognize SLiMs in substrates (11, 12). We recently demonstrated that the disruption of SLiM-substrates interactions could be employed to identify substrates of specific PP2A holoenzyme families (13, 14). However, while SLiM-based substrate recognition seems to be a common mechanism for substrate selection in PPP signaling, SLiMs have only been identified for some PPPs.

Furthermore, SLiMs are commonly shared by regulatory subunit isoforms, not allowing the distinction between isoform-specific functions.

Protein Phosphatase 6 (PP6) is a heterotrimeric holoenzyme, most closely related to PP2A and PP4 (4). The catalytic subunit of the PP6 (PP6c) complex is a highly conserved protein from yeast to humans (4). The PP6 holoenzyme is comprised of a catalytic subunit in a complex with one of three Sit4-associated protein (SAP) subunits, SAPS1, 2, or 3 (also referred to as PP6R1-3), as well as with one of three Ankyrin-domain containing proteins, ANKRD28, 44 or 52 (15, 16). PP6 holoenzymes have been implicated in chromosome condensation and segregation in mitosis (17–23), cell cycle progression (24), inflammatory response (25–28), DNA damage (29–35), splicing (36), maintenance of cell contacts (37, 38), and Hippo (38–40) and mitogen-activated kinase (MAPK) signaling (41). In cancer, PP6C is mutated and deregulated in melanoma (42–47). Hyperactivation of the RAF-MEK-ERK MAPK signaling pathway is frequently observed in cancer, including malignant melanoma, and resistance to drugs targeting kinases in this pathway is common, resulting in therapeutic failure. PP6 was recently shown to be a MEK phosphatase that opposes oncogenic ERK signaling (41). Indeed, loss-of-function PP6c mutations co-occur with BRAF and NRAS mutations in malignant melanoma (41, 44, 45), and loss of PP6c desensitized BRAF-, NRAS-, and KRAS- mutant cell lines to MEK inhibition (41, 48). In addition, the loss of PP6c in flies and mouse keratinocytes promotes RAS-driven tumor growth (49, 50). PP6 was first implicated in Hippo signaling through large-scale proteomics-based interactome studies, which identified a phosphorylation-dependent interaction of Mps One Binder (MOB1) with PP6 holoenzymes upon treatment with the phosphoprotein phosphatase inhibitor okadaic acid (39). MOB1 is a key signal integrator in the Hippo pathway, binding both upstream Ste201Zllike kinases MST1 and MST2, and downstream AGC group kinases LATS1, LATS2, NDR1, and NDR2. MST1/2 bind MOB1 in a phosphorylation-dependent manner and phosphorylate MOB1 on Threonine 12 (T12) and Threonine 35 (T35), inducing a conformational shift that exposes a binding surface for LATS and NDR kinases (51). Long-term depletion of ANKRD28 by siRNA increased MOB1 T35 phosphorylation (52). However, a direct regulatory function of PP6 holoenzymes in the Hippo pathway has not been demonstrated.

The role of PP6 holoenzymes in cellular signaling is complex, and only a few direct dephosphorylation sites have been identified (17, 20, 21, 23, 41). Furthermore, no SLiM has been identified in any PP6 holoenzyme. Because of the lack of specific inhibitors of PPP catalytic subunits, we previously employed RNAi silencing to deplete cells of PP6c and identify PP6c-dependent phosphorylation events (17). Although this strategy successfully identified the condensin I subunit NCAP-G as a PP6 substrate, the prolonged depletion needed for efficient reduction of PP6c expression required orthogonal validation to prove a direct relation. To overcome this limitation, we implemented auxin-inducible protein degradation to rapidly reduce endogenous PP6c levels in near-diploid, KRAS mutant (p.Gly13Asp) DLD-1 cells, followed by mass spectrometry-based quantitative phosphoproteomics to identify and quantify changes in phosphorylation abundances upon auxin (indole-3-acetic acid, IAA) addition and PP6c degradation. The auxin-inducible protein degradation approach combines the cellular ubiquitin-based protein degradation process with a plant-specific F-box protein, Tir1, to induce protein degradation upon treatment with the plant hormone auxin (53, 54). Upon exogenous expression, Tir1 interacts with the cellular Skp1 and Cullin to form a functional SCF E3 ubiquitin ligase. Substrate specificity is dictated by Tir1, which only recognizes its degradation motif in the presence of auxin, resulting in ubiquitination and proteasomal degradation of the auxin-inducible degron (AID) tagged protein (Figure 1A). Using CRISPR-Cas9 genome engineering, any gene can be fused with an AID sequence at the endogenous locus, and the AID-fusion protein is rapidly degraded to study its function. Here, we characterize the AID system to study PP6c function and substrates. We show that within two hours of adding the auxin to mitotic FLAG-sAID-PP6c DLD-1 cells, 95% of PP6c is degraded without any observed off-target effects. We identify significant increases in phosphorylation site abundance on proteins involved in mitotic regulation, organization of the cytoskeleton and cell contacts, and MAPK and Hippo signaling pathways, among others. We validate the direct PP6c-dependent dephosphorylation of MOB1 T35 and demonstrate that PP6c-dependent dephosphorylation regulates the MOB1-LATS1 interaction.

**Figure 1:**
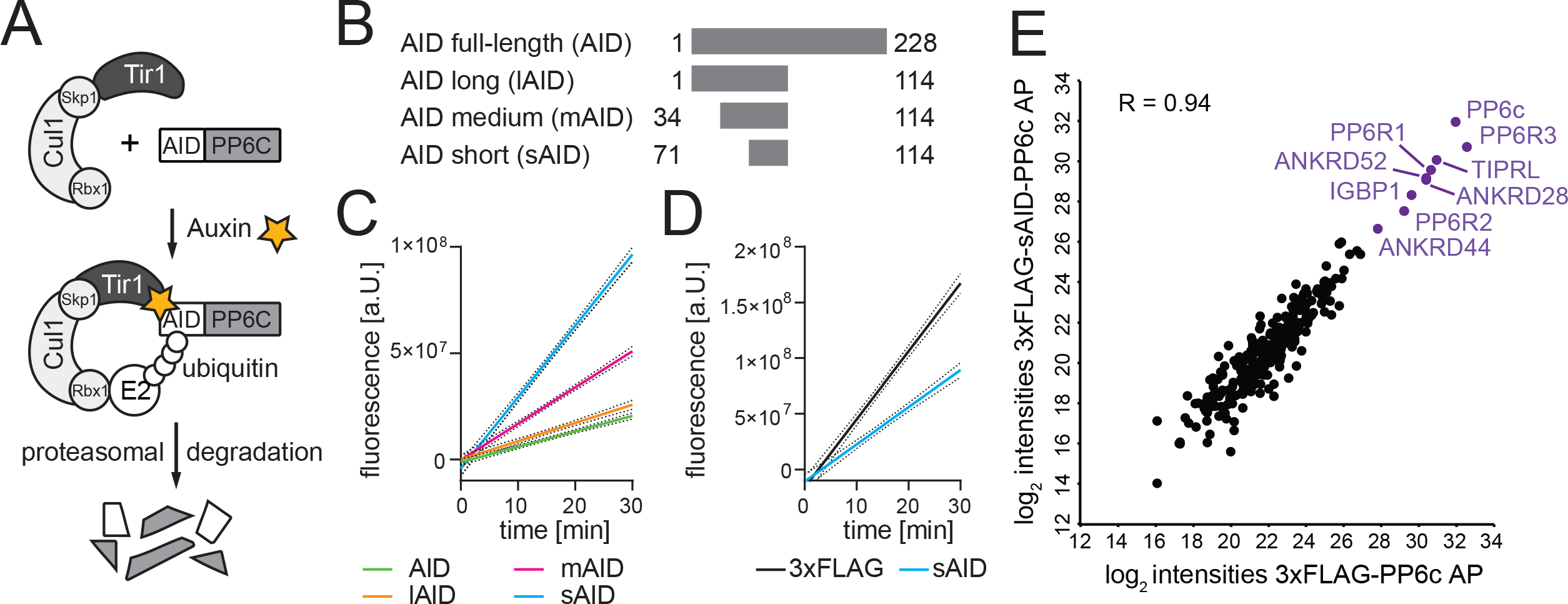
Effects of AID tag variants on PP6c activity in vitro. A) Schematic displaying the mechanism of degradation by the Auxin Inducible Degron (AID) system. Upon the addition of IAA, rapid and targeted proteasomal degradation of the AID-tagged PP6c protein occurs via recruitment of the SCF ubiquitin ligase complex. B) Diagram of all AID tag variants. C) Quantification of phosphatase activity of 3xFLAG-PP6c when fused to the different AID tag variants. D) Quantification of phosphatase activity of 3xFLAG-PP6c compared to 3xFLAG-PP6c fused to the AID short tag variant (sAID). E) Scatter plot of log_2_ protein intensities quantified in 3xFLAG-PP6c and 3xFLAG-sAID- PP6c protein AP-MS experiments. PP6c and known PP6c interactors are shown in purple.

## Experimental Procedures

### Experimental Design and Statistical Rationale

Genome editing was performed using two independent gRNAs predicted by CRISPOR.tefor.net to have limited off-target effects. The sAID was fused to the N-terminus of PPP6c to ensure tagging of the three annotated alternative splice isoforms of PPP6c, and to avoid interference with C-terminal methylation. For proteomic and phosphoproteomic analyses of untreated and auxin (IAA) treated cells, three or four biological replicates were generated and analyzed using our established and reproducible workflow (13, 55, 56). For the proteomic study, 40 μg of peptides were removed from the trypsin digest of each condition. Biochemical and cell biological validation experiments were performed in three biological replicates. Statistical analyses were conducted in Prism (GraphPad) or Perseus (57). Gene ontology analysis and Reactome pathway analyses were performed in Webgestalt (58) and the TOP10 significantly enriched biological processes, cellular components, and molecular functions are reported.

### Cell culture

HEK-293T, HeLa, and DLD-1 cells were grown as adherent cultures, while the HEK293-FT cells were grown as suspension cultures, in Dulbecco’s Modified Eagle’s Medium (DMEM; Gibco) supplemented with 8% heat-inactivated Fetal Bovine Serum (FBS; Hyclone) and 1% penicillin streptomycin (100 U/mL and 100 µg/mL; Corning), at 37°C in a humidified incubator with 5% CO₂.

To synchronize cells in mitosis, 1 mM thymidine (Sigma) was added for 22 hours, followed by a 3-hour washout with PBS (Corning), and subsequent addition of 0.1 μM Taxol (Sigma) for 16 hours. For auxin-induced protein degradation, cells were treated with 1 μM indole-3-acetic acid (IAA) at 37°C for times indicated.

### AID cloning

AID fragment variants were PCR amplified using the following primers: AID Full Length F 5’-CCCAAGCTTATGGGCAGTGTCGAGCTG-3’ AID 1-114 F 5’-CCCAAGCTTATGGCAGTGTCGAGCTGAATC-3’ AID 31-114 F 5’-CCCAAGCTTAGAGGGTTCTCAGAGACGGTTG-3’ AID 71-114 F 5’-CCCAAGCTTAAAGATCCAGCCAAACCTCC-3’ AID R 5’-AAGGAAAAAAGCGGCCGCTACCTTCACGAACG-3’ PCR products were cloned into p3xFLAG-CMV10-PP6c construct (Sigma) using restriction digests and sequence confirmed. Constructs were transfected into HEK-293T cells and selected using G418 to create stably expressing cell lines.

### CRISPR/Cas9 genome editing

For gRNA and Cas9 expression pX330-U6-Chimeric_BB-CBh-hSpCas9 vector (Addgene: Plasmid #42230) and the PPP6c gRNA 2 oligonucleotide were cloned according to Zhang lab protocols and sequence verified (59). gRNAs were selected based upon predicted high-scoring guides from CRISPOR.tefor.net. <u>guide RNAs:</u>

PP6c gRNA 1 5’- ATAACAAGCCGCGGCAACAGCGG - 3’ PP6c gRNA 211 11111111115’- CACGGGCCGGACGTGACGCAGGG - 3’

Homologous recombination (HR) donor constructs were synthesized as gBlocks (IDT) and assembled into the pBluescript II backbone. Homology arms of ∼400-bp spanned the start codon of PPP6c, which included the 5’UTR and the first exon/intron based on the UCSC genome browser genomic sequence. The homology arms were made resistant to all gRNAs used by introducing mutations at the wobble bases of the PAM site and early seed region of the gRNA sequences. Resistance markers (blasticidin or zeocin) followed by a P2A ribosomal skip peptide cleavage site, 3xFLAG, and sAID (71–114) sequences were included.

For the F-box protein expression, a gBlock (IDT) of a codon-optimized version of Oryza sativa Tir1 (Tir1) was synthesized and cloned into a vector containing homology arms targeting the AAVS1 safe harbor site (AAVS1 SA-2A-puro-pA donor vector (Addgene: Plasmid #22075)(60)), a puromycin resistance marker, and a 5xMyc tag. The gRNA_AAVS1-T1 vector (Addgene: Plasmid #41817) was used for targeting the AAVS1 locus (61). For HR at the PPP6c locus, cells were initially transfected using the jetCRISPR (Polyplus) protocol to tag the first PPP6c allele with blasticidin-P2A-3xFLAG-AID and then using the jetPRIME (Polyplus) protocol to tag the second PPP6c allele with zeocin-P2A-3xFLAG-AID. After 48 hours, selection with blasticidin (10μg/mL) or zeocin (100 μg/mL) was started. Selection agents and media were changed every 2 days until cell death halted and colonies formed. Individual colonies were collected and expanded. Codon-optimized Tir1-5xMyc was introduced into the AAVS1 genomic locus of the AID-PP6c DLD-1 clones selected with puromycin (5 µg/ml).

### Screening for AID-PP6c clones

Single-cell clones were allowed to expand to a 48 well plate, then harvested genotyping and western blot analysis. Genomic DNA was isolated using QuickExtract (Epicentre), incubated at 65°C for 6 minutes followed by an incubation at 95°C for 2 minutes using a thermal cycler. For genomic PCR, primers targeting the genomic loci outside of the gBlock’s homology arms and primers internal to each resistance cassette were used to confirm site-specific insertion of resistance cassettes and the loss of wild-type amplicon.

#### Genotyping primers

Forward Genotyping Primer 5’- GCTTAGGCGAGGTTTTCACAG - 3’

New-Scrn-Zeo_R 5’- GAACGGCACTGGTCAACTTGG - 3’

New Scrn-BSD-R 5’- GGTGGATTCTTCTTGAGACAAAGGC - 3’ Reverse Genotyping Primer 5’- GAAGGGGCGAGCCCGCAAATA - 3’

Clones that were positive for tag insertion and loss of wild-type PCR amplicon were expanded and screened for 3xFLAG-sAID-PP6c homozygosity and tag insertion by western blot with PP6c and FLAG antibodies, respectively. Tir1 clones created from homozygous AID clones were screened for Tir1 insertion by western blot with a Myc antibody (9E10, BioXCell). Tir1-positive, sAID-PP6c homozygous clones were finally screened for their capacity to degrade sAID-PP6c by western blot after 4-hour IAA treatment with anti-PP6c and FLAG antibodies.

### Time course treatments

Cells were synchronized and mitotically arrested as described above and treated with 1 μM IAA for 30 minutes, 1 hour, 2 hours, 4 hours, and 6 hours. Cells were then collected, washed with PBS, pelleted, lysed in SDS sample buffer (75 mM Tris-HCl pH 6.8, 2 % SDS, 20 % Glycerol, 0.02 % Bromphenol blue and 100 mM dithiothreitol (DTT)), and boiled at 95°C for 5 minutes, then separated by SDS-PAGE gel electrophoresis. PP6c degradation rate and efficiency were analyzed via western blot. Each PP6c band intensity was quantified (in biological triplicates) using ImageJ and plotted as abundance on a line graph.

### Proteomics and phosphoproteomics TMT experiments

Cells were plated, mitotically arrested, treated with 1 μM IAA, wash with PBS, and harvested in triplicate as described above at the time points indicated. Frozen cell pellets were partially thawed on ice and resuspended in lysis buffer (8 M urea, 50 mM Tris pH 8.1, 150 mM sodium chloride, 2 mM β glycerophosphate, 2 mM sodium fluoride, 2 mM sodium molybdate, 2 mM sodium orthovanadate and EDTA-free Mini-Complete protease inhibitors). Lysates were sonicated three times at 15% power for 10 seconds each on ice, and the protein concentration of the lysates were determined by BCA protein assay. Proteins were reduced with 5 mM DTT at 55°C for 30 minutes, cooled to room temperature, and alkylated with 15 mM iodoacetamide at room temperature for 1 hour in the dark. The alkylation reaction was quenched by the addition of another 5 mM DTT. The lysates were diluted 6-fold in 25 mM Tris pH 8.1, and sequencing grade trypsin (Promega) was added (1:100 w/w) to each lysate and incubated for 16 hours at 37°C. The digests were acidified to pH ≤ 3 by addition of trifluoroacetic acid (TFA; Honeywell), and any resulting precipitates were removed by centrifugation at 7100 RCF for 15 minutes. The acidified lysates were desalted using a 60 mg Oasis desalting plate (Waters). For the proteomic study, 40 μg of each eluate was vacuum centrifuged to dryness and proceeded straight to TMT labeling.

For the phosphoproteomic study, the eluates were vacuum centrifuged for 30 minutes at 37°C to evaporate organic solvent, snap frozen in liquid nitrogen, and lyophilized to dryness. Phosphopeptides were enriched using the High-Select Fe-NTA phosphopeptide enrichment kit protocol, after which the eluates were vacuum centrifuged to dryness, desalted on a 10 mg Oasis desalting plate, and vacuum centrifuged to dryness again.

Both the peptide and phosphopeptide samples were resuspended in 166 mM EPPS pH 8.5, and their assigned TMT reagents, resuspended in acetonitrile (ACN), were added to each sample, vortexed and incubated at room temperature for 1 hour. TMT reporter reagents were assigned to samples as follows. <u>Proteomic</u>: Controls: 126, 128N, 129C; IAA treated for 2 hours: – 127N, 128C, 130N, 131C; IAA treated for 4 hours: 127C, 129N, 130C. <u>Phosphoproteomic</u>: Controls: 126, 127N, 127C, 128N; IAA treated for 2 hours: 128C, 129N, 129C, 130N; IAA treated for 4 hours: 130N, 131N, 131C.

To verify efficient labeling, 5% of each labeling reaction was quenched with hydroxylamine for 10 minutes at room temperature, mixed, and the multiplexed mixture acidified with TFA to pH < 3 before desalting by STAGE tip. Once the labeling efficiency was confirmed to be >95%, the remaining reactions were quenched, combined, and acidified before desalting on an OASIS 10 mg desalting plate. Eluents were dried by vacuum centrifugation before off-line fractionation by HPLC on a pentafluorophenyl (PFP) column (62). The 48 fractions from the HPLC fractionation were concatenated into 16 fractions for the proteomic multiplex and into 24 fractions for the phosphoproteomic multiplex and analyzed by LC-MS/MS on an Orbitrap Fusion Lumos Tribrid instrument or Orbitrap Fusion Tribrid instrument, respectively.

### LC-MS/MS analyses

The phosphoprotoemic analysis was performed on an Orbitrap Fusion Tribrid mass spectrometer (ThermoFisher Scientific, San Jose, CA) equipped with an EASY-nLC 1000 ultra-high pressure liquid chromatograph (ThermoFisher Scientific, Waltham, MA). Phosphopeptides were dissolved in loading buffer (7% methanol, 1.5 % formic acid) and injected directly onto an in-house pulled, polymer-coated, fritless, fused silica analytical resolving column (35 cm length, 100µm inner diameter; PolyMicro) packed with ReproSil, C18 AQ 1.9 µm 120 Å pore stationary phase particles (Dr. Maisch). Samples were separated with a 120-minute gradient of 4 to 33% LC-MS buffer B (LC-MS buffer A: 0.125% formic acid, 3% ACN; LC-MS buffer B: 0.125% formic acid, 95% ACN) at a flow rate of 330 nl/minute. The Orbitrap Fusion was operated in data-dependent, SPS-MS^3^ quantification mode (63, 64), wherein an Orbitrap MS1 scan was taken at 120K resolution and an AGC target value of 3e^5^. The maximum injection time was 100 milliseconds, the scan range was 350 to 1500 m/z, and the dynamic exclusion window was 15 seconds (+/- 15 ppm from precursor ion m/z). Followed by data-dependent Orbitrap trap MS2 scans on the most abundant precursors for 3 seconds. Ion selection; charge state = 2: minimum intensity 2e5, precursor selection range 650-1200 m/z; charge state 3: minimum intensity 3e^5^, precursor selection range 525-1200 m/z; charge state 4 and 5: minimum intensity 5e5). Quadrupole isolation = 0.7 m/z, R = 30K, AGC target = 5e4, max ion injection time = 80ms, CID collision energy = 32%). Orbitrap MS3 scans for quantification (R = 50K, AGC target = 5e4, max ion injection time = 100ms, HCD collision energy = 55%, scan range = 110 – 500 m/z, synchronous precursors selected = 5). The proteomic analysis was performed on an Orbitrap Lumos mass spectrometer operating in data-dependent, SPS-MS^3^ quantification mode wherein an Orbitrap MS1 scan was taken (scan range = 350 – 1400 m/z, R = 120K, AGC target = 2.5e5, max ion injection time = 50 ms), followed by data-dependent ion trap MS2 scans on the most abundant precursors for 1.8 seconds. Ion selection; charge state = 2: minimum intensity 2e5, precursor selection range 600-1400 m/z; charge state 3-5: minimum intensity 4e5). Quadrupole isolation = 0.8 m/z, CID collision energy = 35%, CID activation time = 10 ms, activation Q = 0.25, scan range mode = m/z normal, ion trap scan rate = rapid, AGC target = 4e3, max ion injection time = 40 ms). Orbitrap MS3 scans for quantification (R = 50K, AGC target = 5e4, max ion injection time = 100 ms, HCD collision energy = 65%, scan range = 100 – 500 m/z, synchronous precursors selected = 10).

### Peptide spectral matching and bioinformatics

Peak lists were generated using in-house developed software Rthur 1.0. Raw data were searched using COMET (65) against a target-decoy version of the human (Homo sapiens) proteome sequence database (UniProt; downloaded 2018; 20,241 total proteins) with a precursor mass tolerance of +/- 1.00 Da and requiring fully tryptic peptides with up to 3 missed cleavages, a static mass of 229.162932 Da on peptide N-termini and lysines and 57.02146 Da on cysteines for carbamidomethylation, and a variable mass of 15.99491 Da on methionines for oxidation. For the phosphoproteomic analysis, a variable mass addition of 79.96633 Da on serines, threonines and tyrosine was included. Phosphorylation of serine, threonine and tyrosine were searched with up to 3 variable modifications per peptide and probability of phosphorylation site localization was determined by PhosphoRS (66). The resulting peptide spectral matches were filtered to ≤1% false discovery rate (67). Statistical analyses was performed in Perseus (57). Gene ontology analysis and Reactome pathway analyses were performed in Webgestalt (58).

### Transfection of exogenously expressed proteins

For the exogenous expression of the PP6 holoenzyme, HEK293-FT cells were transfected with pEXPR- IBA105-PPP6c, p3xFLAG-CMV10-3xFLAG-SAPS2, and p3xFLAG-CMV10-Ankrin52 using PEI. For the exogenous expression of the PP6c AID variants, p3xFLAG-CMV10-pFLAG-AID-PP6c vectors were transfected into HEK-293T transiently using PEI and collected after 48 hours. For the generation of stable cell lines, cells were seeded in a 6-well tissue culture dish at ∼50% confluency and transfected using using the JetPrime reagent (Polyplus) per manufacturer’s protocol. After 12 hours, the transfection mixture was replaced with fresh media, and cells were allowed to recover for 24 hours before replating into media containing the respective antibiotics, followed by replenishing the selection agents and media every 2 days until cell death halted and colonies formed. Specifically, for the exogenous expression of Twin-Strep-tagged MOB1, HEK-293T and AID-PP6c DLD-1 cells were transfected with 2µg of pEXPR-IBA105-hMOB1a.

### Affinity purification of twin-Strep-tagged proteins

For the purification of phosphorylated MOB1, HEK-293T cells stably expressing Twin-Strep-tagged hMOB1a were treated with 1 µM calyculin A for 30 minutes and then harvested. For the purification of the PP6 holoenzyme, HEK293-FT cells stably expressing PP6c, PP6R2, and ANKRD52 were harvested. Pellets were washed once with PBS and then lysed in high-salt Strep-Tactin XT buffer (100 mM Tris/HCl, pH 8; 500 mM NaCl; 1 mM EDTA) to a protein concentration of 2 mg/mL and Strep-Tactin XT 4Flow was added to the lysate. Samples were rotated at 4°C for 1 hour, washed four times with the high-salt Strep-Tactin XT buffer, and eluted in 50 mM Biotin dissolved in the high-salt Strep-Tactin XT buffer.

### Affinity purification followed by MS

For the purification of PP6c AID variants, HEK-293T cells transiently expressing 3xFLAG-tagged PP6 variants were lysed in FLAG lysis buffer (50 mM Tris-HCl pH 7.5, 150 mM NaCl, 1% Triton-X100 and a protease inhibitor tablet). Samples were sonicated three times for fifteen seconds each and clarified at 8000 x g at 4°C for 15 minutes. Anti-FLAG M2 affinity gel (SIGMA) previously washed in FLAG lysis buffer was applied to the clarified lysate for 2 hours with rotation at 4°C. Bound resin was then washed three times with FLAG lysis buffer and eluted in 30 µl TBS with 0.16 µg/µl 3xFLAG peptide by vortexing gently for 30 minutes at 4°C. Eluates were then trichloroacetic acid (TCA) precipitated. Briefly, samples were diluted in 20% TCA, incubated on ice for 15 minutes, and centrifuged at 21100 x g for 15 minutes at 4°C. Samples were air-dried, digested with trypsin, and processed for LC-MS/MS analysis.

### PP6 in vitro phosphatase assay

PP6c-AID variant proteins, PP6 holoenzyme trimer and phosphorylated human MOB1a were expressed and purified as described above. For DiFMUP dephosphorylation reactions, purified PP6c AID variants were incubated in reaction buffer (3 mM HEPES pH 7, 0.5 mM DTT, 0.005% Triton-X100, 0.05 mg/ml BSA, 1 mM sodium ascorbate, 1mM MnCl_2_) with substrate 6,8-difluoro-4-methylumbelliferyl phosphate (DiFMUP) and analyzed by fluorescence absorption reading using a Molecular Devices SpectraMax i3x plate reader at excitation 360 nm, emission 450 nm every minute for a 30-minute reaction at 37°C.

For in vitro dephosphorylation reactions, phosphorylated MOB1a and PP6 holoenzyme were combined in the presence or absence of 1 µM calyculin A in phosphatase buffer (30 mM HEPES pH 7, 150 mM NaCl, 0.1 mg/mL BSA, 0.5% Triton X-100, 1 mM ascorbate, 1 mM MnCl_2_, 2 mM DTT) and incubated for 30 minutes at 30°C shaking at 1000rpm. Reactions were quenched by the addition of SDS sample buffer and boiled at 95°C for 5 minutes, then separated by SDS-PAGE gel electrophoresis and analyzed via western blot.

### MOB1 immunoprecipitation

AID DLD-1 cells expressing Twin-strep-MOB1 were arrested in mitosis and IAA treated for 2 hours, as described above. At time of collection, cells were spun down, washed with PBS, and crosslinked with 1% formaldehyde for 10 minutes at room temperature, quenched with 125 mM glycine and washed with cold PBS once more. Cell pellets were lysed in 1.2 mL cold immunoprecipitation buffer (150mM Tris pH 8, 150 mM NaCl, 0.5% Triton X-100, 2 mM sodium fluoride, 2 mM sodium molybdate, 5 mM β glycerophosphate, and EDTA-free Mini-Complete protease inhibitors) and sonicated at 15% power for 10 seconds three separate times. Lysates were clarified by centrifugation (21,000 x g) for 10 minutes at 4°C, and the supernatant incubated with Strep-Tactin XT 4Flow for 1 hour with rotation at 4°C. Immunoprecipitates were washed four times with ice-cold lysis buffer. All proteins were eluted from Strep-Tactin XT 4Flow by addition of SDS sample buffer and boiled at 95°C for 5 minutes, then separated by SDS-PAGE gel electrophoresis and analyzed via western blot.

## Results

### Effects of AID-tags on PP6c activity and holoenzyme formation

To develop an AID strategy for investigating PP6c function, we assessed the impact of fusing an AID tag to the amino (N)-terminus of a PP6c cDNA. We chose N-terminal tagging, because PP6c is modified at its carboxyl (C)-terminus by methyl-esterification (68), and although the function of this modification is still elusive, we wanted to avoid any disruption of PP6c biology by C-terminal tagging. Different length AID tags have been described, all conserving the critical binding domain of Tir1 that confers full degradation capability (Figure 1B) (69, 70). To determine if any of these tags would affect PP6c function, we generated a series of fusion constructs with the different length AID tags. The AID tag variants were amplified by PCR and cloned into an expression vector containing a 3xFLAG-tagged PP6c, resulting in N-terminally tagged 3xFLAG-AID-PP6c fusion products. These constructs, as well as a 3xFLAG-PP6c and 3xFLAG only control were separately transfected into HEK-293T cells and the expressed proteins were purified using the 3xFLAG tag (Supp. Figure 1). The purified 3xFLAG-AID-PP6c variants were tested for enzymatic activity in biological triplicates, using a phosphatase assay which measures the continuous dephosphorylation of a substrate molecule, DIFMUP (6,8-difluoro-4-methylumbelliferyl phosphate), into its fluorescent derivative. This assay revealed that the addition of the AID tag negatively affected PP6c phosphatase activity as a function of length when controlled for total protein amounts (Figure 1C). Particularly, the inclusion of the full-length (amino acids 1-228) and long (lAID, amino acids 1-114) AID tags strongly reduced the phosphatase activity of PP6c to near-baseline levels. PP6c activity was partially recovered upon fusion with the medium length AID tag (mAID, amino acids 34-114). However, the greatest improvement was observed when fusing the shortest AID tag length (sAID, amino acids 71-114) to PP6c, which recovered ∼61% of the 3xFLAG-PP6c control activity (Figure 1D).

To further validate that the 3xFLAG-sAID-PP6c fusion protein functionally compared to the 3xFLAG-PP6c, we performed affinity purification followed by mass spectrometry (AP-MS) analysis to identify potential differences in holoenzyme assembly and protein-protein interactions. We purified 3xFLAG-sAID-PP6c and 3xFLAG-PP6c, TCA precipitated elutes, digested proteins into peptides with trypsin, and analyzed them by LC-MS/MS. This analysis revealed a high correlation (R = 0.94) between the proteins that co-eluted from each condition, with the PP6 regulatory subunits PP6R1-3, ANKRD28, ANKRD44, ANKRD52, and the known PP6c interactors IGBP1 and TIPRL being highly enriched and their abundances correlated between both proteins (Figure 1E). Thus, although the N-terminal tagging of PP6c with the shortest AID-tag partially reduced PP6c phosphatase activity, it preserved the overall holoenzyme composition and protein-protein interactions of PP6c.

### Generation of 3xFLAG-sAID-PP6c DLD-1 cells using CRISPR/Cas9-based endogenous tagging

To investigate endogenous PP6c function in cells using the AID strategy, we utilized CRISPR/Cas9- mediated homology-directed repair to introduce a 3xFLAG-sAID tag into the genomic locus of PPP6c in DLD-1 cells. We chose DLD-1 cells because they are near-diploid and are shown to be chromosomally stable, which meant only two rounds of endogenous targeting would be necessary to tag all PPP6c alleles (71). There are three annotated alternative splice isoforms of PPP6c, all of which begin translation at Exon 1’s starting methionine (Supp. Figure 2A). We reasoned that insertion of the AID sequence at the N-terminus of PPP6c should result in the tagging of all expressed splice isoforms. To introduce the AID system, we designed a targeting construct for homologous recombination (HR) in a pBlueScript II backbone containing: a resistance cassette with alternate selection markers to be driven by the PPP6c promotor, a ribosomal skip sequence (P2A peptide cleavage site), a 3xFLAG tag, and the sAID (71–114) degron sequence, all in frame with the PPP6c ATG start codon. These elements were flanked by PPP6c homology arms, corresponding to 400 bp directly upstream and downstream of the transcription start site, which contained silent mutations against the gRNA target sites to protect the recombined alleles from further editing (Figure 2A and Supp. Figure 2B). DLD-1 cells were co-transfected with the gRNA/Cas9-containing pX330 dual expression vector and the blasticidin-specific PP6c target construct and placed under antibiotic selection. Single colonies were expanded and screened by genotyping and western blot analysis. Once a clone was proven to have a successful insertion, the process was repeated with a zeocin resistance marker containing targeting construct.

**Figure 2:**
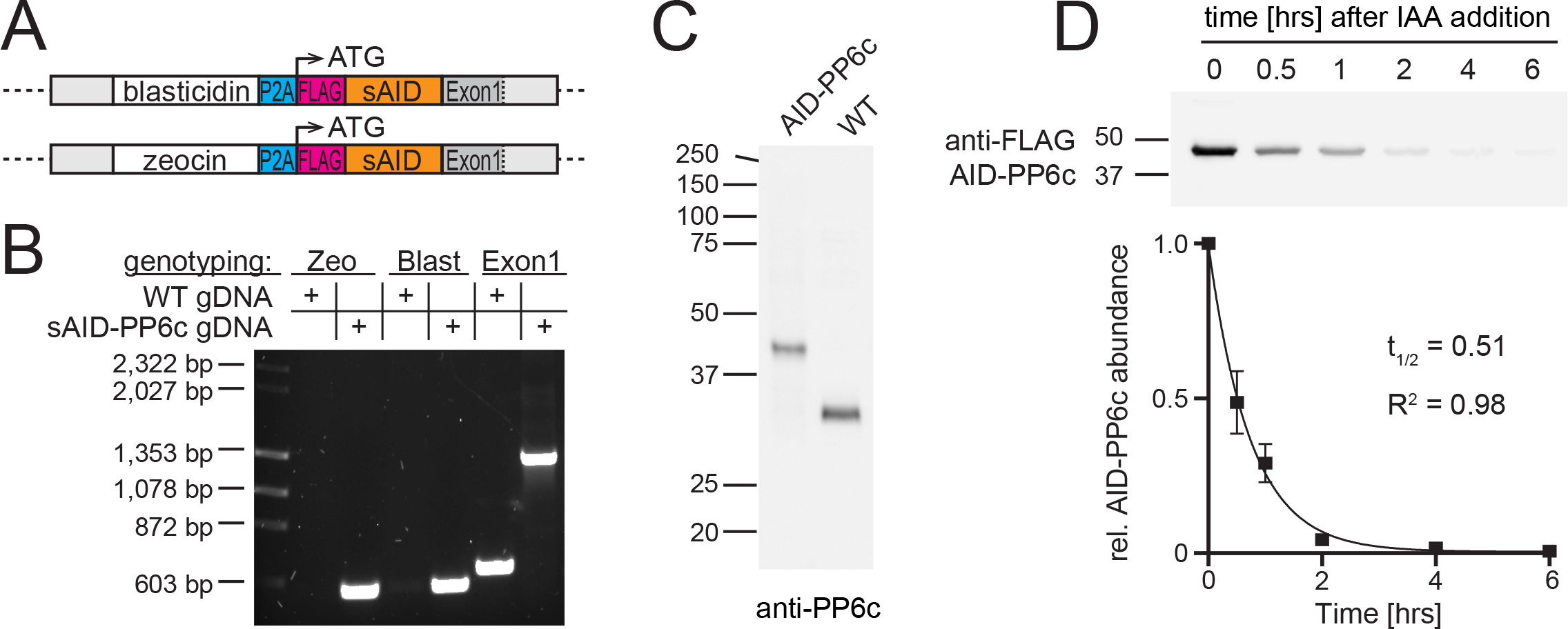
Endogenous tagging of PPP6c genomic loci in DLD-1 cells. A) Schematic depicting the PPP6c genomic alleles in DLD-1 cells upon homologous recombination with the targeting constructs. From left to right: left homology arm (light grey) consisting of the 5’UTR, a resistance marker (white), P2A ribosomal skip site (cyan), 3xFLAG tag (magenta), the sAID tag (orange), and the right homology arm consisting of Exon 1 (dark grey) and the beginning of Intron 1 (light grey). B) Agarose gel analysis of genomic PCRs for the PPP6c loci in wild-type (WT) and homozygously tagged 3x- FLAG-sAID-PP6c DLD-1 cells. We will refer to these cells from here on as AID-PP6c DLD-1s and to the tagged protein as AID-PP6c. C) Western blot analysis of endogenous PP6c protein in WT and AID-PP6c DLD-1 cells. D) Top: Western blot analysis of time course experiment demonstrating inducible degradation of AID-PP6c protein upon addition of IAA. Bottom: Quantification of AID-PP6c protein abundance overtime. Data are presented as means11±11standard deviations from three independent experiments. The AID-PP6c protein half-life was calculated by fitting exponential regressions to the averages from triplicate quantifications.

For genotyping, we chose a forward primer that annealed outside of the left homology arm, and reverse primers directed towards the blasticidin or zeocin resistance marker resulting in 564 bp or 556 bp amplicons, respectively (Supp. Figure 2B and Figure 2B). Because DLD-1 cells are diploid, the presence of both resistance markers indicates that both PPP6c alleles were tagged. To further demonstrate that no wild-type (WT) PPP6c alleles remained, we performed an additional genotyping PCR with a reverse primer that annealed in Exon 1, resulting in a PCR product of ∼629 bp for WT alleles or ∼1,322 bp or ∼1,298bp for alleles that included the inserted degron and the blasticidin or zeocin resistance cassette, respectively (Supp. Figure 2B). Because of the similar size of the recombined alleles (1,322 bp and 1,298 bp), the two PCR products migrate closely together and are not separated by gel electrophoresis (Supp. Figure 2B). Furthermore, we performed western blotting using an antibody against endogenous PP6c. Only clones that displayed a complete band shift of endogenous PP6c protein, from the WT molecular weight of 35 kDa to the molecular weight of the fusion protein at 42 kDa, were chosen for further analysis. A representative western blot displaying a complete band shift is seen in Figure 2C, indicating the fusion of sAID and FLAG tags to all endogenously expressed PP6c protein, hereafter referred to as AID-PP6c. Finally, we performed western blot analysis with an anti-FLAG antibody to confirm the inclusion of 3xFLAG epitope tag (Figure 2D).

To induce ubiquitylation and subsequent proteosomal degradation of PP6c upon auxin addition, we introduced the auxin-dependent F-box protein, Tir1, into the AAVS1 safe harbor locus of the homozygously-tagged 3xFLAG-sAID-PP6c DLD-1 cell line. For this, we used a targeting construct for homologous recombination consisting of a puromycin resistance marker followed by the Tir1 coding sequence fused to a 5xMyc tag and flanked by homologous regions to the AAVS1 locus. The targeting construct was co-transfected with a px330 dual expression vector containing Cas9/gRNA cDNA and cells were selected with puromycin (61). After single-cell cloning, resistant clones were screened by western blotting with an anti-Myc antibody to detect Tir1 expression (Supp. Figure 3). To determine if Tir1 was functional, Tir1 positive clones were treated with 1 µM of auxin (IAA) and probed for auxin-dependent degradation of PP6c. This yielded several clones with functional Tir1 that degraded PP6c upon the addition of IAA. Finally, to identify a clone with fast degradation kinetics, we performed a time course analysis of the IAA-dependent degradation of AID-PP6c, choosing a clonal cell line that exhibited efficient degradation in two hours (Figure 2D). We will refer to this cell line hereafter as AID-PP6c DLD-1 cells.

**Figure 3:**
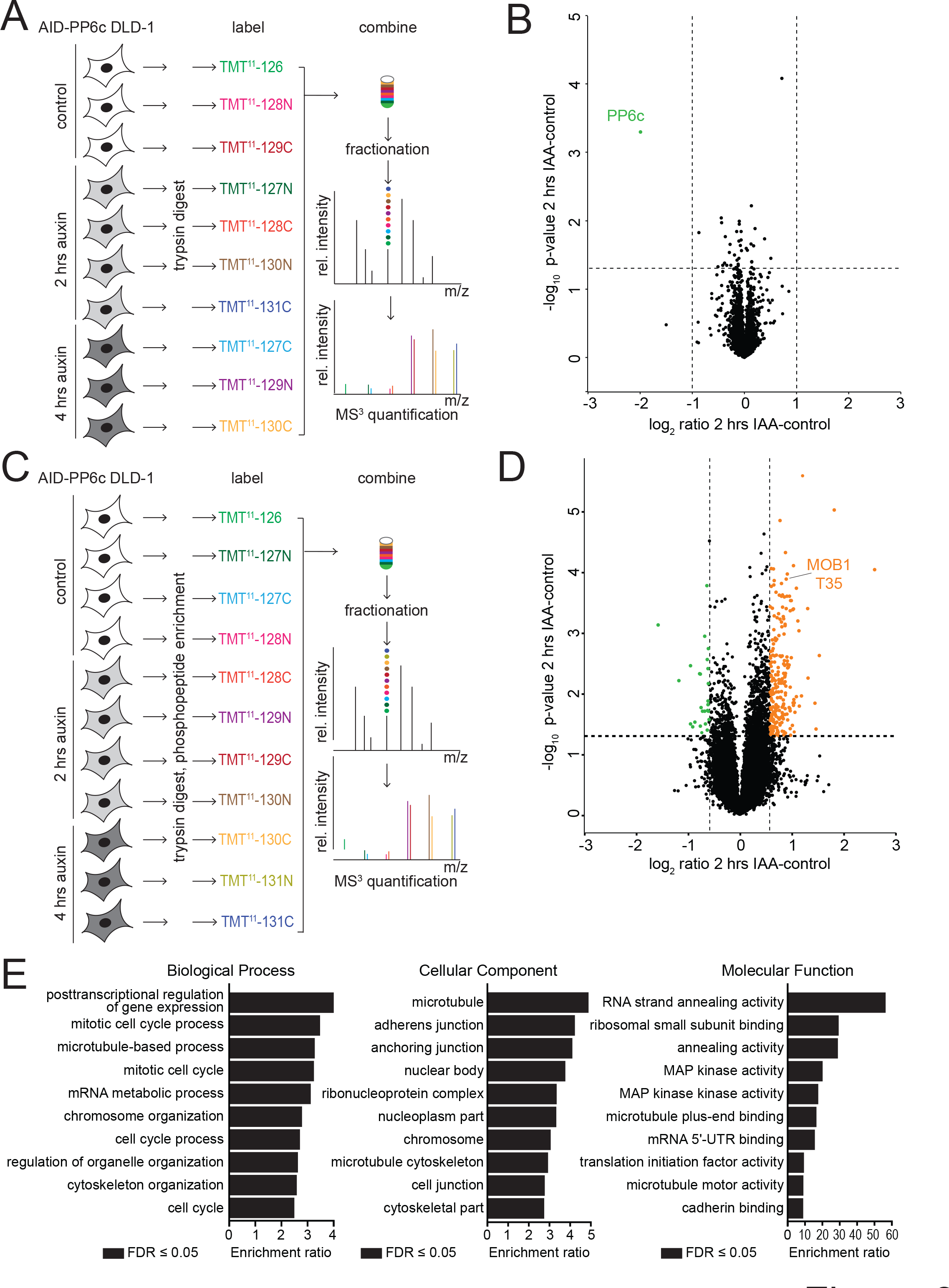
Quantitative proteomics and phosphoproteomics reveal potential PP6c substrate dephosphorylation sites. A) Workflow for TMT multiplex proteomic analysis of mitotic AID-PP6c DLD-1 cells wherein, three channels were assigned to control conditions (n=3), four channels to 2 hours IAA treatment conditions (n=4), and three channels to 4 hours IAA treatment conditions (n=3). B) Volcano plot of the log_2_ ratio fold-change of proteins identified and quantified in the 2-hour IAA treated versus untreated cells. The vertical lines represent fold-change cutoffs of -1 and 1, the horizontal line represents a significance cutoff of a p-value of 0.05, and each dot represents one protein. PP6c is indicated in green. C) Workflow for TMT multiplex phosphoproteomic analysis of mitotic AID-PP6c DLD-1 cells wherein four channels were assigned to control conditions (n=4), four channels to 2 hours IAA treatment conditions (n=4), and three channels to 4 hours IAA treatment conditions (n=3). D) Volcano plot, showing the log_2_ ratio fold-change of phosphorylation sites identified and quantified in the 2 hours IAA treated versus untreated cells. The vertical lines represent fold-change cutoffs of -0.6 and 0.6, the horizontal line represents a significance cutoff of a p-value of 0.05, and each dot represents a phosphorylation site. Significantly increased phosphorylation sites are highlighted in orange, while decreased sites are shown in green E) Top 10 categories from gene ontology analysis of proteins with significantly upregulated phosphorylation sites as input.

### IAA-inducible degradation is highly selective for AID-PP6c protein in mitotically-arrested DLD-1 cells

Next, we investigated the selectivity of IAA-dependent protein degradation in the AID-PP6c DLD-1 cell line. We performed global quantitative proteomics of untreated and IAA-treated cells. In brief, cells were synchronized in mitosis with a thymidine block followed by Taxol arrest and then either treated with 1 µM IAA for 2 hours or 4 hours or not treated in biological replicates. After collection, cells were lysed, proteins were digested into peptides using trypsin, and peptides were labeled with tandem mass tag (TMT) reagents to enable multiplexing of all conditions and replicates. Samples were pooled, and peptides were fractionated before LC-MS^3^ analysis (Figure 3A). We identified and quantified 7,484 proteins in this experiment, of which 6,007 had a total peptide (TP) count of two or more. Indeed, we found that the only protein with a significant fold-change of greater than two upon IAA addition was PP6c (Figure 3B). Overall, these data demonstrate the successful introduction of the AID system into DLD-1 cells and that auxin-inducible degradation is an effective and selective method to reduce specific protein abundances in cells with no observable off-target effects at our detection level.

### Quantitative phosphoproteomics analysis of mitotically-arrested, IAA-treated AID-PP6c DLD-1 cells

Having confirmed the selectivity and specificity of the IAA-inducible depletion of PP6c, we investigated changes in the phosphoproteome of mitotically-arrested AID-PP6c DLD-1 cells upon 2 hours and 4 hours of IAA treatment. Again, we followed the same procedure as described previously for the proteomics analysis with the addition of a phosphopeptide enrichment step after the trypsin digestion (Figure 3C). Here we identified 23,278 phosphorylation sites on 5,526 proteins, of which 288 and 57 were significantly increased or decreased by 1.5-fold or more, respectively (Figure 3D). We observed a high correlation (R = 0.85) between the significantly quantified phosphorylation sites in the 2-hour and 4- hour treated conditions (Supp. Figure 4A).

**Figure 4:**
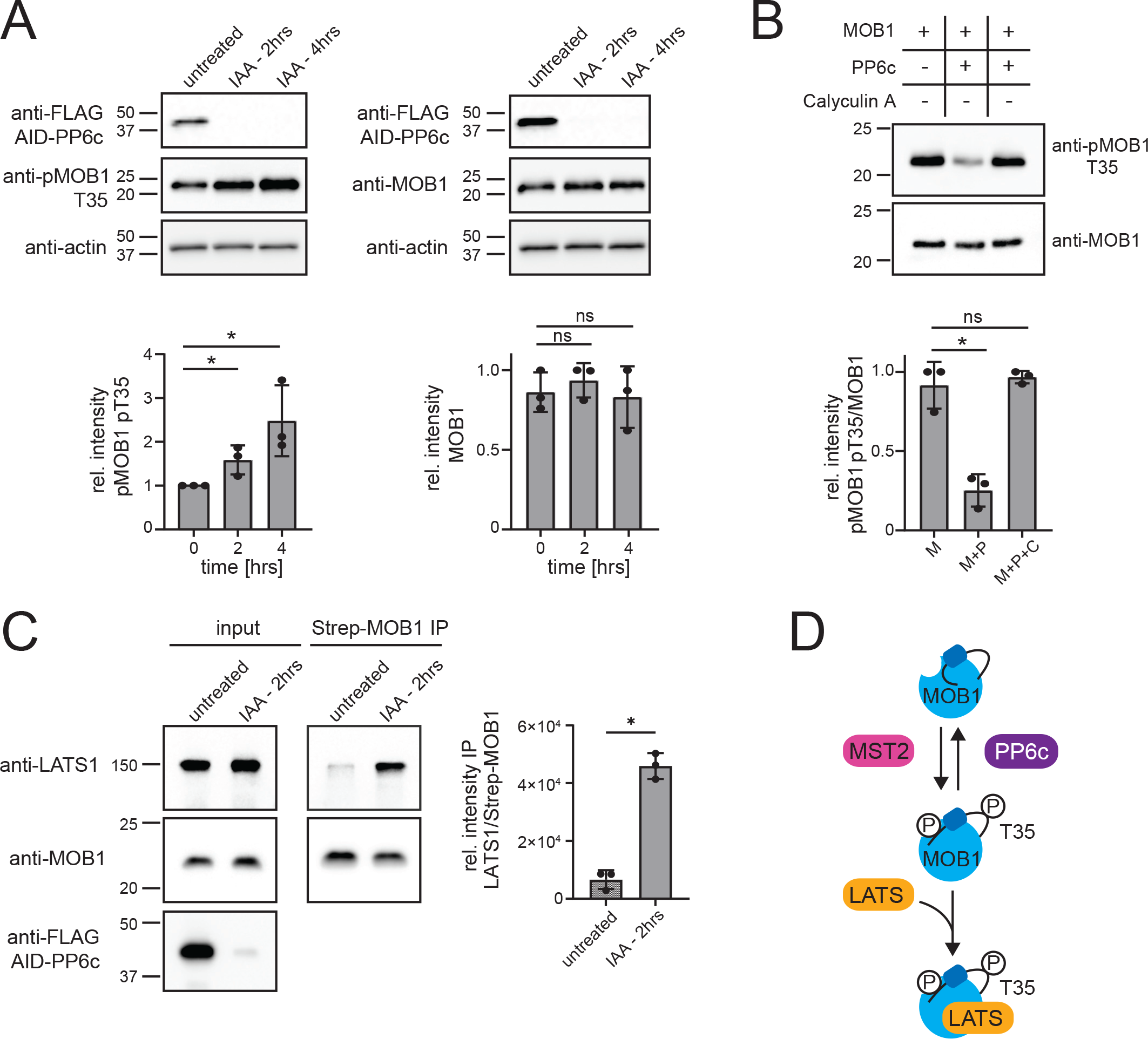
MOB1 T35 phosphorylation is regulated by PP6c activity. A) Top left: Western blot analysis of the abundance of AID-PP6c protein and MOB1 T35 phosphorylation in untreated versus IAA-treated AID-PP6c DLD-1 cells, with β-actin used as a loading control. Bottom left: Quantification of MOB1 T35 phosphorylation relative to the β-actin control. Top right: Western blot analysis of the abundance of AID-PP6c and MOB1 protein in untreated versus IAA-treated AID-PP6c DLD- 1 cells, with β-actin used as a loading control, using the same lysates as on the left. Bottom right: Quantification of MOB1 total protein relative to the β-actin control. For both bar graphs, data are presented as means11±11SD using lysates from the same three independent experiments, and individual data points are indicated. *p < 0.05, ns – not significant by Student’s t-test. B) Top: Western blot analysis of an in vitro dephosphorylation assay using affinity-isolated PP6c holoenzyme incubated with affinity- purified phosphorylated MOB1 (pMOB1) in either the presence or absence of 1 µM calyculin A inhibitor (CalA). Bottom: Quantification of MOB1 T35 phosphorylation relative to the level of MOB1 total protein. On the x-axis: “M” is pMOB1 only, “M+P” is pMOB1 combined with active PP6 holoenzyme and “M+P+C” is pMOB1 combined with PP6 plus the addition of CalA. Data are presented as means11±11SD from three independent experiments, and individual data points are indicated. *p < 0.05, ns – not significant by Fisher’s exact t-test. C) Left: Western blot analysis of input and affinity-purified MOB1 and its downstream interactor LATS1 in untreated compared to IAA treated AID-PP6c DLD-1 cells exogenously expressing twin-Strep-tagged MOB1. Right: Quantification of LATS1 abundance relative to the amount of MOB1 that was pulled down. Data are presented as means11±11SD from three independent experiments, and individual data points are indicated. *p < 0.05, ns – not significant by Student t-test. D) Representative diagram showing a proposed model for PP6c’s negative regulation of the Hippo Pathway’s core signaling cascade.

To broadly investigate the list of phosphorylation sites that were significantly increased by the rapid degradation of PP6c, we performed gene ontology (GO) term and pathway enrichment analyses using the Reactome database. The GO terms revealed that our dataset is enriched in biological processes linked to mitotic cell cycle regulation, chromosome organization, and posttranscriptional regulation of gene expression (Figure 3E). Likewise, the pathway analysis paralleled these findings with the identification of enrichment in pathways connected to mitotic spindle checkpoint and chromosome cohesion signaling, as well as mitotic cell cycle phases such as prometaphase, metaphase, and anaphase (Supp. Figure 4B). PP6c is well known for its role in mitotic regulation through direct dephosphorylation of mitotic substrates such as the Condensin complex I, and also indirectly through the regulation of Aurora kinase A (Aurora A) (17, 20, 23). PP6 directly dephosphorylates the Aurora A activation T-loop at Threonine 288 (T288), with previous reports detailing how the reduction of PP6 activity can lead to the hyperphosphorylation of Aurora A substrates (17, 20). Indeed, we observed increased phosphorylation of Tastin Serine 156 (S156) and NDC80 Serine 55 (S55) sites, both of which are known Aurora A substrates (72, 73).

In terms of cellular components, adherens, anchoring, and cell junctions were among the most enriched terms, along with cadherin binding emerging as a highly enriched term in the molecular function category (Figure 3E). Thus far, limited studies have been done to connect PP6 to cell adhesion regulation. Intriguingly, one report found that PP6 localizes to adherens junctions and opposes Casein kinase 1 to maintain E-cadherin levels at the cell surface. Moreover, the study found that PP6c protein abundance appears to be regulated by cell density in multiple epithelial cell lines (37). In our analyses, the over-representation of cell adhesion molecule binding and extracellular matrix constituents represented on the list of PP6c substrates suggests strong evidence for a new and yet-unexplored role of PP6c in regulating cell adhesions and different types of cell junctions.

Among the most upregulated phosphorylation sites were the Tyrosine 204 (Y204) and Tyrosine 187 (Y187) sites of ERK1 and ERK2, respectively (Supp. Figure 4A). Although PP6c does not have any known tyrosine phosphatase activity, Cho et al (41) reported activation of the RAF-MEK-ERK pathway upon deletion of PP6c due to its acting as a direct phosphatase and inhibitor of ERK’s upstream kinase MEK. Our analyses agree with these findings as the observed activation of the MAP kinase signaling pathway, demonstrated by ERK hyperphosphorylation, was also prominent in the molecular function and pathway enrichment analyses (Figure 3E, Supp. Figure 3).

Much like the RAF-MEK-ERK pathway, the Rho family-GTPases are a subgroup of the Ras superfamily of GTP-binding proteins that function as signal transduction molecules in the cell. Intriguingly, the phosphoproteomics, and subsequent bioinformatic analyses, strongly implicate PP6 in the regulation of the Rho signaling pathway, evidenced by the fact that signaling by Rho GTPases and Rho GTPase effectors are among the highest represented terms in the pathway analysis (Supp. Figure 3). It is well-known that the primary function of the Rho pathway is to mediate the dynamic rearrangements of the plasma membrane-associated actin cytoskeleton, which is also reflected in the GO term analysis result that indicated cytoskeleton organization and cytoskeleton part as significantly enriched terms in the molecular function and cellular components categories, respectively. To our knowledge, there are no studies that have identified a role for PP6c in the context of Rho signaling. However, a study using a colorectal cell line with the same RAS activating mutation as DLD-1 cells discovered that PP6c co-precipitated with RhoA (74). Furthermore, in two separate interactome studies, PP6 subunits were found to interact with ARHGEF2, a direct binder and regulator of the RhoA protein kinase (39, 75). Furthermore, several proteins with significantly increased phosphorylation sites upon PP6c degradation are either GEF activators of the RhoA protein kinase or known substrates. One of these sites is Rho-associated coiled–coil domain-containing protein kinase 2 (ROCK2) S1372, which is phosphorylated in mitosis by Polo-like kinase 1 (Plk1) (76). This is intriguing as it suggests that through its action on Aurora A, PP6c indirectly regulates Plk1. Aurora A is the activating kinase of Plk1 (77, 78), and inhibition of PP6c activity appears to result not only in the activation of Aurora A but also Plk1 (23). Thus, our data suggest direct and indirect hyperactivation of the Rho/ROCK kinase cascade upon PP6c depletion. Overall, our observations confirm our approach’s validity by identifying known substrates and pathways regulated by PP6c, and they reveal novel regulatory roles of PP6c that are still yet to be examined.

### Investigation of PP6c-dependent MOB1 signaling

MOB1 is an integral component of the Hippo pathway, serving as the scaffold protein responsible for assembling the core kinase cascade that propagates Hippo signaling. MOB1 is phosphorylated by mammalian Ste20-like kinases MST1 and MST2 at several residues at its N-terminus, including Threonine 35 (T35), and these phosphorylations induce a conformational shift that exposes a binding surface for the downstream kinase, LATS1 (39, 40). Multiple PP6 subunits were previously found to interact with MOB1 in AP-MS and proximity labeling experiments (39, 40). This discovery then led to additional biochemical studies that were able to independently validate the direct interaction between MOB1 and PP6 protein (39, 40).

In our phosphoproteomic analysis, we identified MOB1 S9 and T35 as phosphorylation sites that were significantly increased upon degradation of PP6c (Supp. Table 3). To validate this finding, we performed western blot analysis in biological triplicates using a phospho-site-specific antibody against the MOB1 T35 site and confirmed a significant increase in T35 phosphorylation upon PP6c degradation while MOB1 protein levels remained unchanged (Figure 4A). To determine if there is a direct regulatory function linking MOB1 and PP6c, we performed an in vitro phosphatase assay using purified PP6 holoenzyme (consisting of PP6c, PP6R2, and ANKRD28) and purified phosphorylated MOB1 protein. MOB1 T35 was readily dephosphorylated upon incubation with PP6 holoenzyme, and this activity was blocked by the addition of the PPP inhibitor calyculin A (Figure 4B). Since the phosphorylation of MOB1 T35 promotes the interaction of LATS1 (51), we reasoned that PP6c degradation, and subsequent increase in phosphorylation of MOB1 T35, should enhance the MOB1-LATS association. To assess whether there was a change to MOB1 interaction with LATS1 upon PP6c degradation, we stably expressed exogenous Twin-Strep-MOB1 protein in the AID-PP6c DLD-1 cell line. Cells were arrested in mitosis, treated with IAA for 2 hours, and Twin-Strep-MOB1 was affinity enriched. The elutes were analyzed by probing the MOB1 pulldowns for associated LATS1 via western blot analysis. This revealed a significant increase in LATS1 binding to MOB1 upon PP6c degradation (Figure 4C). Taken together, these results support the notion that a PP6 holoenzyme opposes MST1/2 phosphorylation at T35 on MOB1, limiting the interaction of MOB1 with LATS1 and thereby the activation of the core Hippo kinase cascade (Figure 4D).

## Discussion

Reversible protein phosphorylation is an essential regulatory mechanism controlling most cellular processes. The phosphorylation state of a protein substrate depends on the opposing activities of protein kinases and protein phosphatases. Advances in methodology and instrumentation of mass spectrometry-based phosphoproteomics have greatly increased our understanding of the regulatory role of protein kinases and their specific substrate phosphorylation sites (2, 79). However, compared to their counteracting kinases, our knowledge of the opposing protein phosphatases is often limited. This is partly due to the lack of tools and strategies for studying phosphatases, especially serine/threonine phosphatases. (80). In eukaryotes, the PPP family of phosphatases is responsible for the majority of serine/threonine dephosphorylation in the cell (4). The small number of PPP catalytic subunits achieve substrate specificity by forming multimeric holoenzymes with hundreds of regulatory subunits. However, the highly conserved and shallow nature of the active site of the PPP catalytic subunits hampers specific targeting attempts (6, 7). Thus, to elucidate PPP signaling, alternative approaches are needed.

In our previous work, we used shRNA depletion of the PP6 catalytic subunit to identify regulated signaling pathways, which found Condensin I as a PP6 substrate and a role of PP6 in chromosome condensation (17). While successful, this approach is hampered by the need for prolonged depletion to ensure sufficient reduction in the catalytic subunit abundance, making it difficult to distinguish between direct and indirect effects. Therefore, as an alternative approach, we sought to establish a strategy for a more direct method of PPP substrate identification capable of rapid depletion of all cellular PP6 activity. Thus, we turned to developing an auxin-inducible degradation approach to investigate PP6 signaling.

The original AID-tag adds ∼25 kDa to the target protein (Figure 1B) and, like other larger tags, can affect protein function. To determine the effect of the AID-tag on PP6c, we made N-terminal fusion protein containing a 3xFLAG- and AID-tag, expressed it in HEK-239T cells, and tested the purified protein by in vitro phosphatase assay. Compared to the activity of 3xFLAG-PP6c, the phosphatase activity of the AID-tagged fusion protein was greatly diminished (Figure 1C). We overcame this by reducing the size of the AID tag, which restored phosphatase activity (Figure 1D). This demonstrates that in vitro testing of the enzymatic activity of AID-fusion proteins is essential to ensure biological function.

Because DLD-1 cells are diploid for the PPP6c gene, it was necessary and sufficient to introduce the degron tag into the two PPP6c genomic loci to control all endogenous PP6 holoenzymes and their activity. Furthermore, endogenous tagging of the PPP6c alleles, as opposed to exogenous expression of degron-tagged PP6c, ensured that we would most closely recapitulate PP6 biology regarding subunit composition, abundances, and phosphorylation signaling. Therefore, using homologous recombination and CRISPR/Cas9 gene targeting, we introduced a short version of the AID tag (Figure 1B) into the genomic loci of PPP6c in DLD-1 cells and selected a clone with rapid degradation kinetics (t_1/2_ = 0.51 hrs) (Figure 2D).

Quantitative proteomics analysis of this cell line identified and quantified 7,484 proteins with only the PP6c protein significantly decreasing in abundance upon treatment with IAA. These results demonstrate that the IAA treatment is specific and selective for AID-tags in human cells and produces no detectable off-target effects among the 7,484 proteins quantified. While the untagged proteome remained stable, IAA treatment resulted in an increase of 288 phosphorylation sites. In contrast, only 57 phosphorylation sites were decreased, supporting the notion that the observed increases in phosphorylation are due to ceased PP6c activity upon its targeted degradation.

The phosphorylation sites with the largest observed increases upon IAA treatment compared to control were the kinase activating sites, ERK1 Y204 and ERK2 Y187, while showing no significant change in protein abundance. It was recently shown, using shRNA-mediated depletion and an in vitro phosphatase assay, that PP6 negatively regulates MEK kinase activity by dephosphorylating the activation sites on its T-loop in cell lines with activating RAS mutations, resulting indirectly in the downstream activation of ERK (41). Furthermore, loss of function mutations in PPP6c in KRAS mutant lung and pancreatic ductal adenocarcinoma, and NRAS mutant lung adenocarcinoma and melanoma resulted in resistance to small molecule MEK inhibitor (48). In KRAS mutant mouse keratinocytes and flies, loss of PPP6c promotes tumorigenesis (49, 50). Also, in human melanomas, PPP6c is frequently mutated and these mutations often occur in combination with BRAF or NRAS mutations (44, 45). As DLD-1 cells are heterozygous for a KRAS p.Gly13Asp activating mutation, our observations of MEK hyperactivation upon acute PP6c degradation further supports a direct role for PP6c as a tumor suppressor in a RAS mutant context.

One of the best-characterized functions of PP6 is its regulatory role as the deactivating T-loop phosphatase of Aurora A, an essential mitotic kinase implicated in microtubule spindle formation and chromosome alignment (17, 20, 23, 43). In mitosis, Polo-like kinase 1 phosphorylates a PP6 holoenzyme, inhibiting its activity towards Aurora kinase A (23). Although our analysis did not detect any Aurora kinase A peptides, it did reveal an enrichment in substrates with a function in the mitotic cell cycle, microtubule-based processes, and chromosome organization. Furthermore, we observed increased phosphorylation of Tastin and NDC80 on sites known to be phosphorylated by Aurora A (72, 73). Increased Aurora A activity is observed in human cancer, and selective small molecule inhibitors have been developed and tested in clinical trials (81). Indeed, melanoma cells expressing PP6c mutant proteins are more sensitive to Aurora A inhibition than control cells or cells expressing wild-type PP6c (82).

An emerging function of PP6 is as a negative regulator of the Hippo signaling pathway (39, 40, 52). PP6 subunits interact with MOB1 in a phosphorylation-dependent manner (39, 40), and long-term depletion of ANKRD28 increased MOB1 T35 phosphorylation, regulating focal adhesion dynamics in cell migration (52). Our analysis revealed a significant increase in S9 and T35 MOB1 phosphorylation upon PP6c degradation, as well as an enrichment in the cellular component category for cell junctions. Using an in vitro phosphatase assay, we were able to confirm a direct regulatory role of PP6 in the dephosphorylation of the T35 site of MOB1. Finally, we showed that the degradation of PP6c increased not only MOB1 T35 phosphorylation but also the interaction of MOB1 and the LATS kinases. It was previously shown that phosphorylation of MOB1 on T12 and T35 induces a conformational shift creating a binding surface for LATS and NDR kinases (51). Our demonstration of the direct dephosphorylation of MOB1 T35 by PP6 functionally links the interaction of PP6 and MOB1 with the regulation of the phosphorylation-induced interaction and activation of MOB1 and LATS1 (83, 84). Activated LATS kinases phosphorylate YAP (yes-associated protein) and TAZ (WW domain–containing transcription regulator protein 1), inhibiting their nuclear localization. YAP/TAZ are transcriptional co-regulators that bind to TEAD transcription factors to promote growth, cell proliferation, and cell survival. Activation of Hippo signaling prevents YAP/TAZ-TEAD target gene expression and many Hippo pathway components, including LATS1/2 and MOB1, are characterized as tumor suppressors. Thus, in the context of Hippo signaling, PP6c seems to act as an oncogene whose inactivation results in the activation of the Hippo pathway. This is in contrast to PP6c’s role in MAPK and Aurora A signaling, where the loss of PP6c activity leads to hyperactivation of oncogenic kinase signaling, as described above. These differences might be dependent on the relative expression of the different PP6 regulatory and scaffolding subunits and their assembly into distinct holoenzymes with diverse substrate specificities, cellular localization, and tissue expression. Furthermore, the RAS mutation status and other kinase signaling alterations are likely to impact PP6 function. In the future, it will be important to investigate PP6c substrates in various cancer and normal cell lines and use AID strategies to rapidly deplete regulatory and scaffolding subunits. While long-term depletion by siRNA and gene disruption of one subunit can lead to compensation by other isoforms, rapid protein degradation should overcome this limitation and allow us to decipher subunit-specific signaling pathways.

## Acknowledgments

We thank members of the Gerber and Kettenbach labs for helpful discussion and comments. The work was supported by grant R35GM119455 from NIGMS to ANK and NSF GRFP 1840344 to NCM. Shared resources and the Molecular Genomics Core at the Dartmouth Cancer Center are supported by NIH/NCI P30CA023108. The pX330-U6-Chimeric_BB-CBh-hSpCas9 vector (Addgene: Plasmid #42230) was a gift from Dr. Feng Zhang. The gRNA_AAVS1-T1 vector (Addgene: Plasmid #41817) was a gift from Dr. George Church. The AAVS1 SA-2A-puro-pA donor vector (Addgene: Plasmid #22075) was a gift from Dr. Rudolf Jaenisch.

## Data availability

Mass spectrometry data have been deposited to ProteomeXchange: PXD037202 and MassIVE: https://massive.ucsd.edu/ProteoSAFe/dataset.jsp?task=cc064f9e966e495aaec879c9a20c95ab. Reviewer password: p1293.

---

**Supplemental Figure 1:**
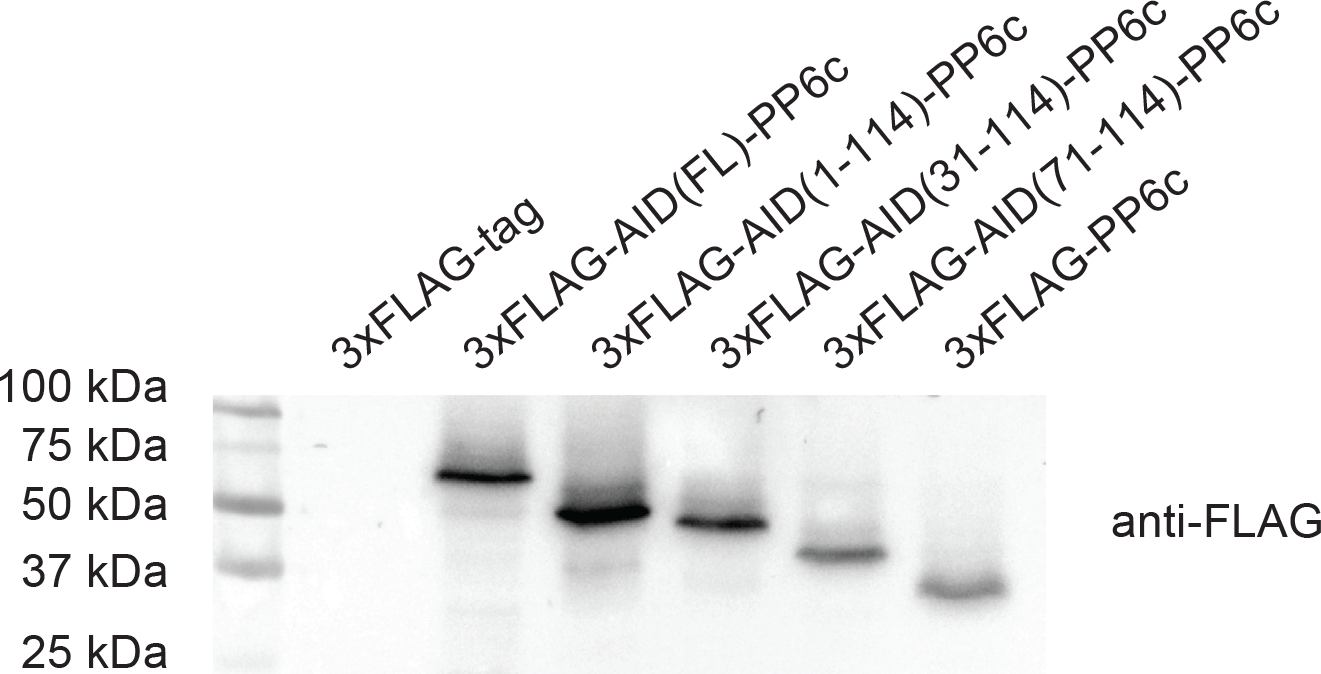
Expression of 3xFLAG-AID-PP6c variants and 3xFLAG-PP6c. Western blot analysis of 3xFLAG-tag, 3xFLAG-AID-PP6c variants, and 3xFLAG-PP6c affinity purifications.

**Supplemental Figure 2:**
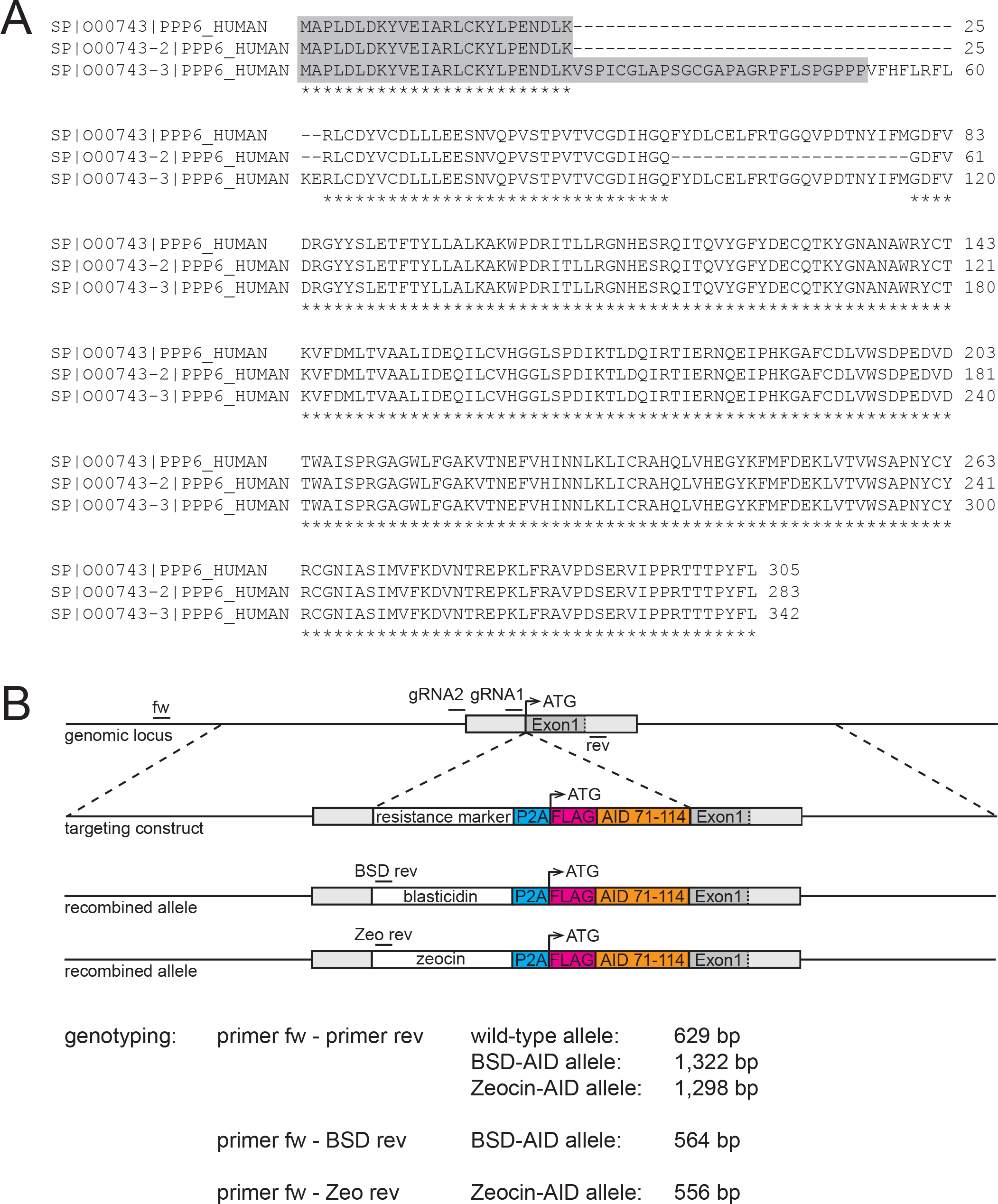
PP6c isoforms and targeting strategy. A) Amino acids sequences of PP6c isoforms. B) Schematics depicting the PPP6c genomic locus with guide RNAs (gRNAs) and genotyping primers (fw and rev) indicated, the targeting construct (from left to right: left homology arm (light grey) consisting of the 5’UTR, a resistance marker (white), P2A ribosomal skip site (cyan), 3xFLAG tag (magenta), the sAID tag (orange), and the right homology arm consisting of Exon 1 (dark grey) and the beginning of Intron 1 (light grey)), and the recombined alleles with blasticidin and zeocin resistance marker and location of their specific genotyping primers indicated. Sizes of the expected PCR products are listed below.

**Supplemental Figure 3:**
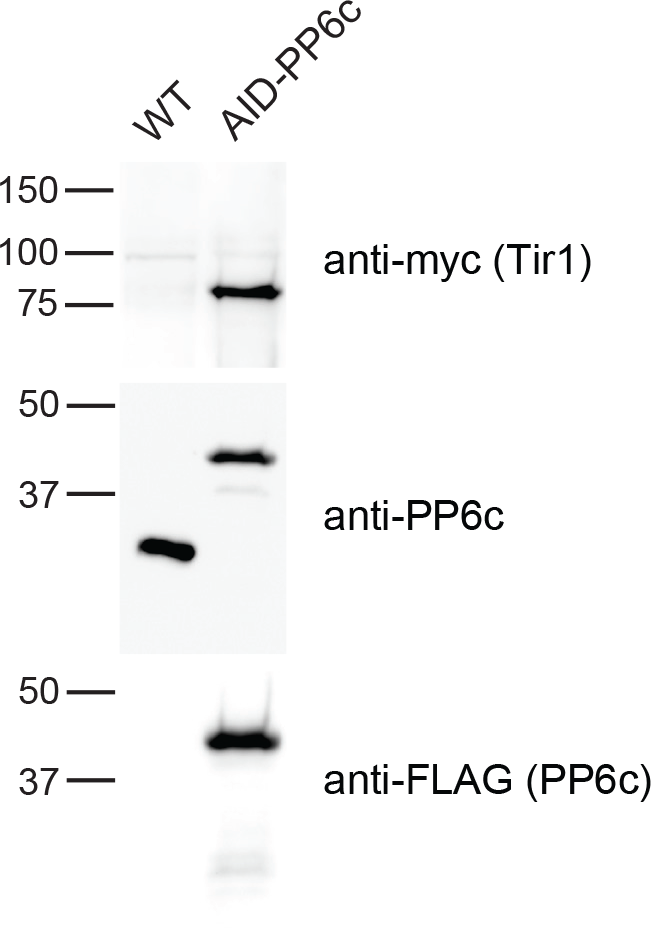
Tir expression in AID-PP6c DLD-1 cells. Western blot analysis of the abundance of myc-Tir1 protein (top), endogenous PP6c (middle), and FLAG-PP6c (bottom) in WT and AID-PP6c DLD-1 cells.

**Supplemental Figure 4:**
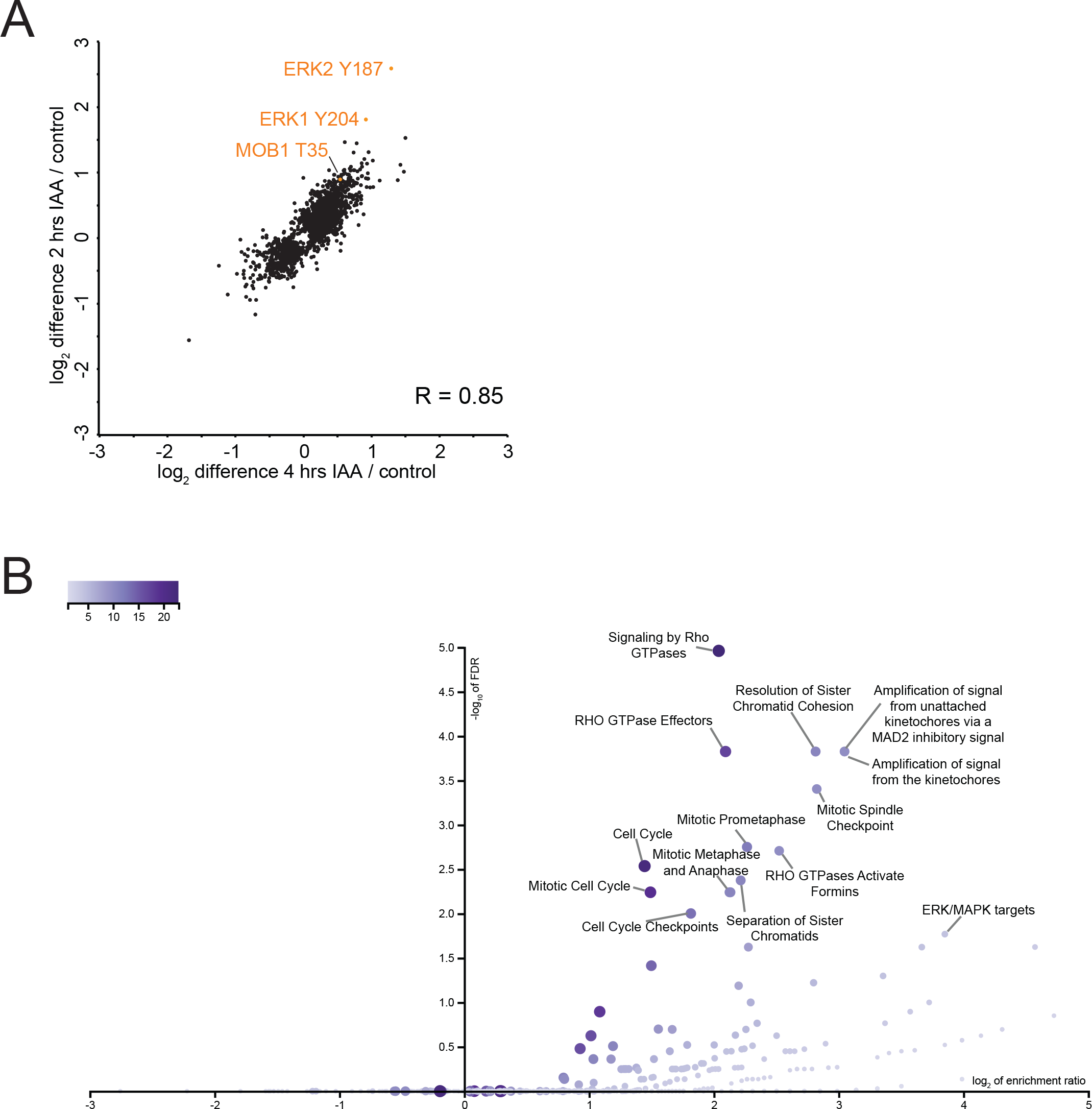
Analysis of AID-PP6c phosphoproteomics dataset. A) Scatter plot of significantly quantified log2 differences of phosphorylation sites in 2 and 4 hours IAA treatment compared to control. ERK1 Y204 and ERK2 Y187, phosphorylation sites known to increase upon PP6c depletion, and MOB1 T35 are indicated in orange. B) Volcano plot of the log2 of enrichement ratio of Reactome pathways containing proteins with significantly increased phosphorylation sites.

